# ERα mediated gene state switching regulates the extent of the single-cell estrogen response

**DOI:** 10.1101/2023.10.18.562977

**Authors:** Christopher R. Day, Pelin Yasar, Gloria Adedoyin, Brian D. Bennett, Joseph Rodriguez

## Abstract

Gene regulation is complex, involving the coordination of hundreds of proteins that function to control genome accessibility, mediate enhancer-promoter interactions, and initiate transcription. At individual loci, transcriptional initiation is stochastic, resulting in short periods of nascent RNA synthesis known as transcriptional bursts. To understand how altered Estrogen Receptor function and cofactor recruitment regulates transcriptional bursting, we used single molecule imaging of estrogen responsive genes in Bisphenol A (BPA) treated cells. Using live cell imaging of the estrogen responsive *TFF1* gene, we observe that cells treated with BPA exhibited burst initiation kinetics and burst sizes which were indistinguishable from cells induced with Estradiol (E2). However, we observed a 50% reduction in the number of active alleles in BPA treated cells. This effect is gene specific, as *GREB1* was unperturbed. Although we observed no difference in chromatin accessibility, the *TFF1* promoter exhibited an altered structure which coincided with reduced ERα and cofactor binding. Lastly, deletion of the enhancer locus removed the BPA effect, indicating that enhancer function was perturbed. Our results demonstrate gene specific effects of altered ERα recruitment and function which lead to a reduction of transcriptionally permissive states. Our work supports the model that the early estrogen response occurs from alleles in primed transcriptionally permissive states with additional inactive alleles contributing to the response over time.

## Introduction

Transcriptional initiation requires the precise coordination of numerous factors with specialized functions. At single alleles, this complex regulation results in stochastic episodes of nascent RNA synthesis known as transcriptional bursts [1, 2]. Bursting is initiated by the recruitment of transcription factors to regulatory elements in the enhancer and promoter [3, 4]. In the hormone response, the Estrogen Receptor α (ERα) is bound by the primary circulating estrogen in the body, 17-β-estradiol (E2)[5]. This binding causes a conformational change in the ligand binding domain which transitions the ERα into an active form [6, 7]. Through specific DNA binding at regulatory elements and cofactor recruitment, the ERα coordinates chromatin remodeling, including Histone 3 Lysine 27 acetylation (K27ac) and chromatin accessibility [8, 9].

In response to increasing E2 levels, transcriptional initiation or burst frequency increase, resulting in higher RNA output [10, 11]. However, under saturating E2 concentrations, only a fraction of alleles is active. Moreover, the *TFF1* gene exhibits highly variable periods of inactivity in between bursts which can last from minutes to several days [10]. Therefore, regulation of estrogen responsive genes in the presence of higher E2 levels must involve increasing the number of alleles that transition into transcriptionally permissive states and increasing burst initiation rates.

The conformational changes observed after E2 binding result in the generation of new interaction surfaces for cofactors at the Ligand Binding Domain (LBD) [12]. Proper cofactor interactions are crucial for transcriptional activation. Therefore, cofactor recruitment dynamics can have a profound impact on transcriptional bursting [13, 14]. Among the families of cofactors crucial to the estrogen response are the p160 proteins, which encompass Steroid Receptor Coactivators (SRC-) 1, 2, and 3. These cofactors play pivotal roles in nuclear receptor mediated transcriptional activation, including ERα signaling [15]. SRC family members are recruited by ligand-bound ERα complexes and enhance transcriptional activity through cooperation with p300 and CARM1 [16]. Another central complex in gene activation is the mediator complex, which regulates interactions between enhancers and promoters. Mediator promotes gene activation by interacting with TFIID, nuclear receptors, and RNA polymerase II (PolII) [17, 18]. MED1, a specific subunit of the mediator complex, was identified as a critical transcriptional coactivator for ERα-mediated transcription facilitating interactions between the nuclear receptor, the core mediator complex and PolII [19].

Notably, the efficiency of ERα interactions with several cofactors including MED1 and SRC-3 is altered when the receptor is bound by estrogen-mimicking chemicals known as endocrine-disrupting chemicals (EDCs) [20, 21]. Humans are continuously exposed to a complex mixture of EDCs including genistein (Gen) from soy products, Bisphenol S (BPS), and Bisphenol A (BPA) from plastics [22, 23]. Treatment with these chemicals elicits reduced ERα binding to chromatin, unstable ERα conformations and an altered transcriptional response [24–26]. However, it is unknown how these effects alter transcriptional bursting.

Here, we use single molecule imaging to understand how altered estrogen receptor function regulates transcriptional bursting of estrogen responsive genes. We find that BPA treatment maximally induces *TFF1* burst initiation, however there is a significantly smaller number of active alleles. We determine that ERα, FOXA1 and MED1 recruitment are reduced at specific ERα target genes including *TFF1*. Deletion of TFF1’s enhancer results in similar levels of transcriptional activation with both E2 and BPA, suggesting that the ERα complex recruited by BPA disrupts enhancer function. Finally, chromatin accessibility at *TFF1* is increased with BPA treatment. However, the *TFF1* promoter adopts an altered structure which positions a nucleosome over the ERα binding site.

## Results

### Fewer cells transcribe *TFF1* alleles in BPA induced cells

Transcriptional activation requires alleles to switch from a transcriptionally nonpermissive state to a permissive state. To investigate how disruption of ERα cofactor recruitment alters this transition between transcriptional states, we used endocrine disruptor chemicals (EDC) to induce ERα-mediated transcriptional bursts. We performed single-molecule fluorescence *in situ* hybridization (smFISH) of the E2-responsive gene *TFF1* (Figure 1A), to measure RNA content and bursting features. We used probe sets targeting the exons of *TFF1* to visualize and quantify RNA. Bound probe sets are visible as discrete fluorescent spots (green). To identify transcription sites (TS) we used probe sets targeting the first *TFF1* intron (red). Nuclear intron spots that colocalized with exon spots are identified as TSs (Figure 1B). We first performed a dose response in MCF-7 cells to determine the doses for maximal induction with each EDC. We treated cells with either E2 or the three EDCs, genistein (Gen), Bisphenol S (BPS) and Bisphenol A (BPA) for 65h to allow the cells to achieve steady state. While *TFF1* RNA per cell saturates at 1nM E2, all EDCs induced *TFF1* RNA at significantly higher doses of 10mM or more (Figure 1C, and 1D). Gen and BPS treatment resulted in similar *TFF1* mRNA per cell as E2 treatment. However, saturating doses of BPA resulted in 45% lower mRNA per cell (Figure 1E). We also determined the fraction of cells actively transcribing *TFF1* by extracting the percentage of cells with 1 or more TS and observed on average 19.6% of E2 treated cells had active *TFF1* transcription. Gen and BPS treatment also resulted in a similar fraction of cells actively transcribing *TFF1* compared to E2 treatment. However, in the presence of BPA we observed only 9.0% of cells actively transcribing *TFF1* (Figure 1F). We next extracted the number of RNA produced at the TS by normalizing the intensity of the TS by the median signal of the cytoplasmic RNA [10]. We observe a slight increase in nascent *TFF1* transcripts at the TS in cells treated with Gen relative to E2 (p=0.02), but no changes in nascent *TFF1* transcripts in cells induced with BPS or BPA (Figure G). Finally, we performed a time course with saturating levels (1nM for E2 or 10mM for BPA were used for the rest of the assays) of either E2 or BPA to determine whether the difference in *TFF1* RNA accumulation with BPA treatment occurred rapidly after treatment. The time course revealed that both ligands induce transcription at similar rates within the initial two hours (h) of treatment. However, we observed that there is a difference that is maintained after 4h of induction (Figure 1H). These data indicate that Gen and BPS treatment induces *TFF1* bursts which are similar to E2 induction. However, active *TFF1* transcription is reduced by BPA treatment when compared to E2, resulting in 50% less cells with active TS and lower RNA accumulation.

**Figure 1:**
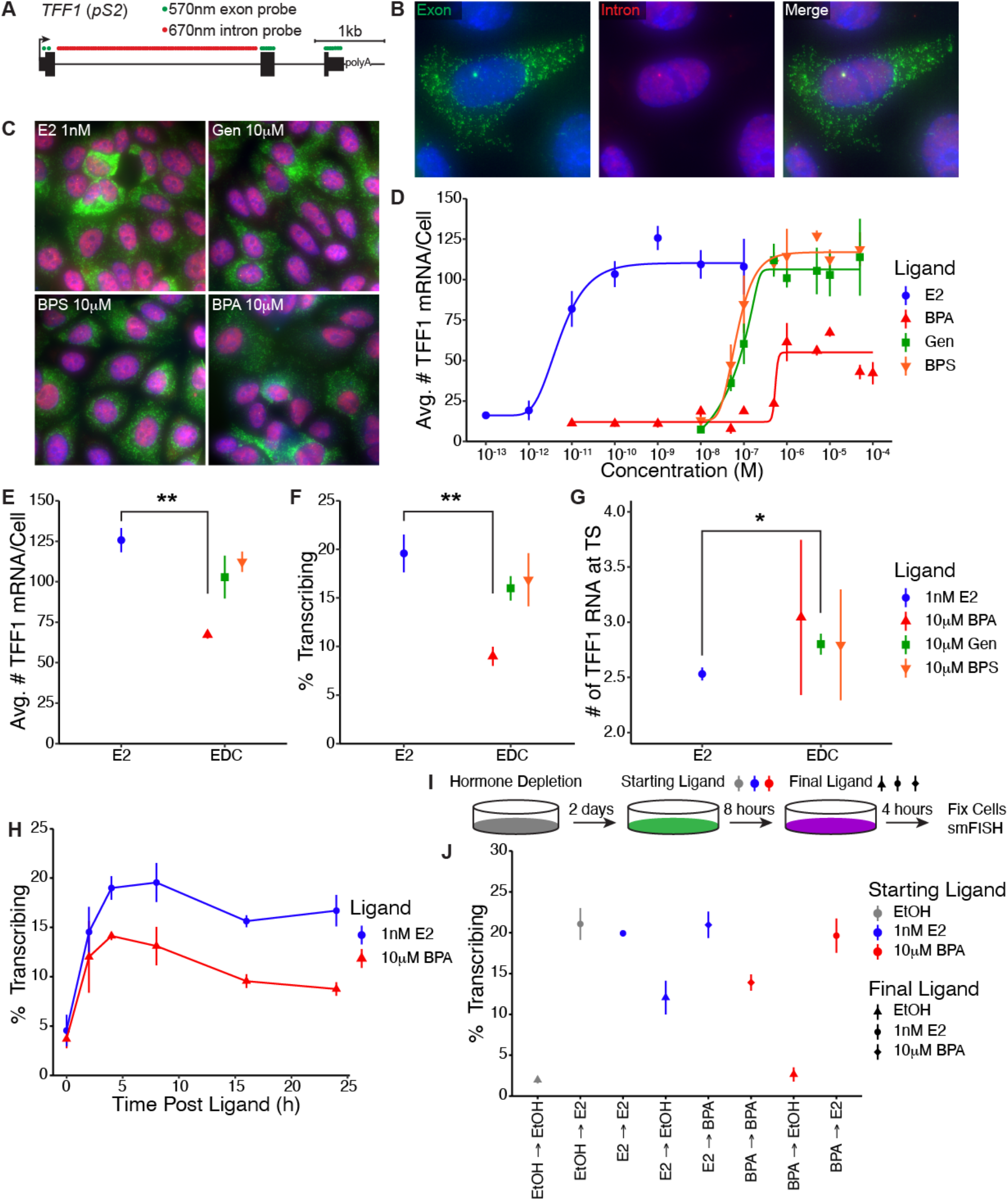
A. A schematic detailing of smFISH probes specific to *TFF1* gene exons and intron. B. A representative image of *TFF1* smFISH, DAPI (Blue), cytoplasmic *TFF1* mRNA (green) and intron (red). TS visible in the nucleus where exon and intron probes colocalize. C. *TFF1* induction was demonstrated in response to 1 nM E2, 10 µM Gen, 10 µM BPS, or 10 µM BPA. D. Dose dependent accumulation of TFF1 mRNA was calculated based on smFISH results in response to E2, BPA, Gen, or BPS. E. Average number of *TFF1* mRNA per cell. Error bars represent standard deviation of the average number of *TFF1* mRNA per cell from 3 replicates. P-values were calculated with t-tests. F. Average percent of cells actively transcribing TFF1 from 3 replicates. P-values were calculated with t-tests. G. The number of TFF1 RNA at TS were determined through smFISH in response to saturating levels of E2 or EDCs. Statistical differences between E2 and EDCs were calculated with t-tests. H. The percentage of actively transcribing cells was plotted over time (0-25 hours) in response to saturating levels if E2 or BPA. I. An illustration depicting the experimental design for a washout experiment. J. Percentage of actively transcribing cells were calculated for each condition and statistical differences were determined in between conditions.

To determine if the BPA effects on bursting were reversible, we performed a washout experiment. Cells were hormone depleted similarly to previous experiments before induction with either saturating levels of E2, BPA or vehicle for 8h. This initial induction followed by a washout with the reciprocal compound or vehicle for additional 4h (Figure 1I and J). We then performed smFISH to determine the fraction of cells actively transcribing *TFF1*. In cells initially treated with vehicle, 4h of E2 was sufficient to significantly upregulate *TFF1* transcription resulting in 21.07% of cells transcribing *TFF1* (Figure 1J, gray markers) again indicating that transcription of *TFF1* is maximally induced after 4h of E2. In cells initially induced with E2, a 4h wash out and replacement with vehicle resulted in a half maximal response in the fraction of cells transcribing *TFF1*, indicating that E2 was not completely washed out. However, pretreatment with E2 followed by a wash out with BPA rescued the activation and 20.96% of cells actively transcribing *TFF1* (Figure 1J, blue diamond). Conversely BPA pretreatment and replacement with vehicle results in a similar fraction of cells transcribing *TFF1* as in vehicle only treated cells (Figure 1J, triangles). Pretreatment with BPA, and replacement with E2 lead to a higher fraction of cells with transcription sites and a comparable response to E2 only treatment. Taken together these results indicate that the BPA perturbation on *TFF1* transcription is reversible, and that BPA primarily activates E2-primed alleles.

### Bursting dynamics are similar between E2 and BPA induced alleles but fewer alleles are active

Our smFISH results indicate that fewer cells have *TFF1* transcription sites when induced with BPA. This data suggests that *TFF1* transcription dynamics are altered in one of two ways. First the inactive periods between bursts could be longer in cells induced with BPA, resulting in less frequent *TFF1* transcriptional bursts across the cell population. Alternatively, fewer *TFF1* alleles could be in a transcriptionally permissive state in the population. To distinguish between these two mechanisms, we used an MCF-7 cell line where 3 *TFF1* alleles are tagged at the endogenous locus with 24xMS2 loops in the 3’ UTR (Figure 2A). The MS2-eGFP recognizes and binds the stem-loops formed by the nascent RNA, coating the transcript with up to 48 fluorophores [27]. Thus, active transcription sites are visible as a diffraction-limited spot and can be used to measure bursting dynamics over time (Figure 2B). *TFF1* bursting dynamics were extracted from *TFF1-MS2* cells that were induced with saturating concentrations of E2 or BPA at least 8h prior to imaging. We tracked individual *TFF1-MS2* alleles to quantify burst and inactive state durations [10, 28]. Bursting alleles were observed in cells induced with both E2 and BPA. Furthermore, the extracted fluorescence intensity traces revealed similar numbers of bursts between cells induced with E2 and BPA, with alleles in both conditions bursting only once and other alleles bursting more frequently in 14h. (Figure 2C, Movie 1 and 2). We observed a small decrease in median burst duration in cells induced with BPA (8.33min with E2 v 6.67min with BPA, Figure 2D). However, our smFISH data indicates the shorter burst duration does not lead to fewer RNA produced per burst (Figure 1G). Surprisingly, we observed no difference in the median inactive times between E2 and BPA (50min v 51.67min respectively, p=0.238, Figure 2E). These data indicate that BPA maximally induces *TFF1* burst frequency relative to E2 induction.

**Figure 2:**
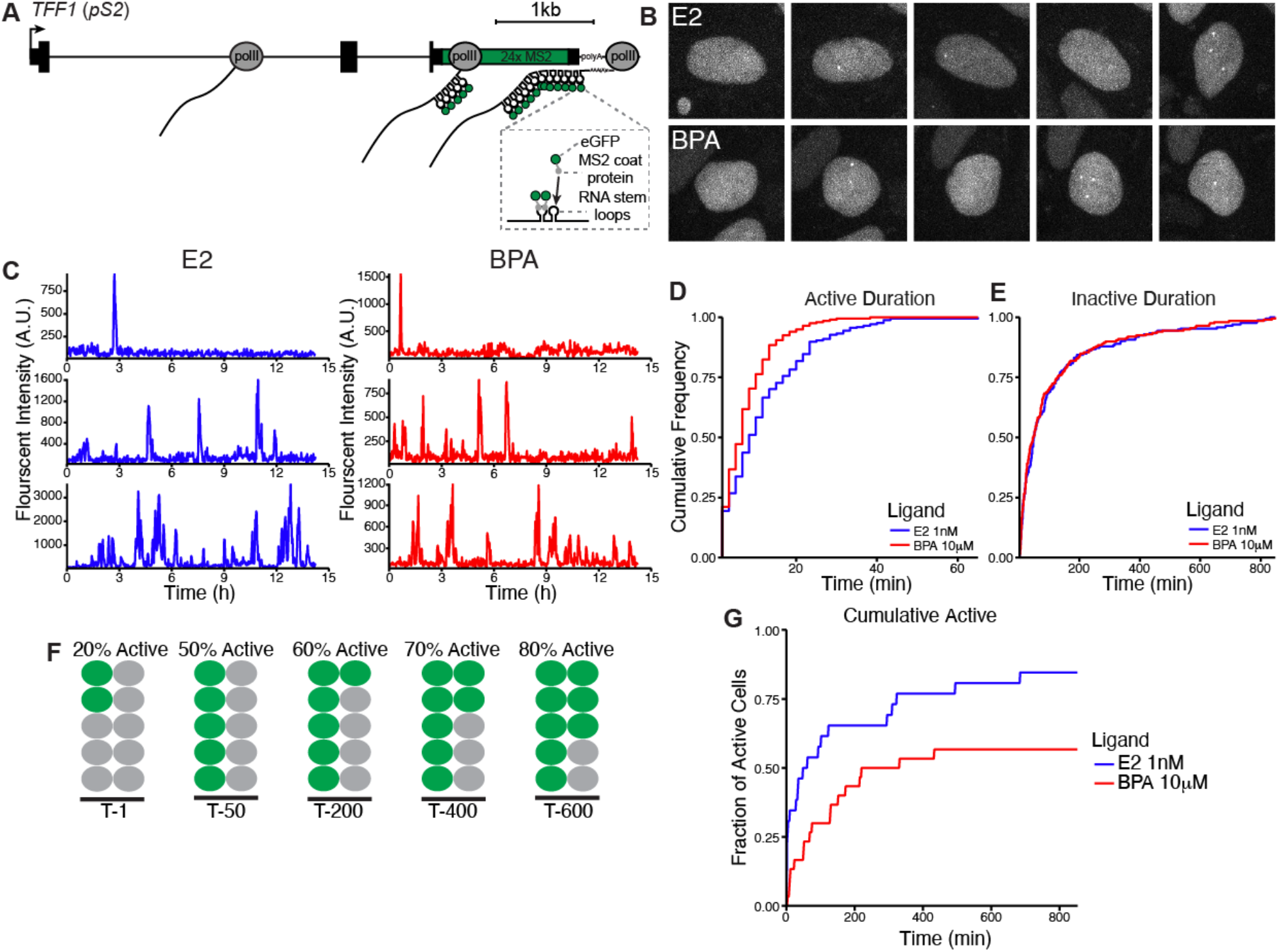
A. A schematic of *TFF1* alleles labeled with 24xMS2 stem loops. B. Visualization of active *TFF1-MS2* transcription in cells induced with E2 and BPA. C. Multiple tracked alleles from cells induced with E2 and BPA. D. Cumulative distribution functions (CDF) of the active durations in cells induced with E2 (blue) and BPA (red). E. Cumulative distribution of the inactive durations between bursts with E2 (blue) and BPA (red). F. Representation of the fraction of active cells. G. Cumulative distribution of the faction of active cells induced with E2 and BPA.

Our data indicate that fewer alleles are transcriptionally active in the population with BPA treatment. To verify this observation we calculated the cumulative fraction of cells that exhibited a transcription site over time. Nuclei were extracted from our imaging data and a machine learning approach was used to automatically segment transcription sites in the nuclei. The cumulative fraction of active cells was determined by dividing the number of nuclei with transcription sites by the total number of nuclei as a function of time (Figure 2F). We observe that a smaller fraction of cells exhibited a *TFF1* burst over the duration of the time course. Moreover, there is a 25% difference in the fraction of cells with *TFF1* bursts in BPA induced cells relative to E2 induced cells, which plateaus for the BPA treatment after 6.5h of imaging (Figure 2G). Taken together our single-molecule imaging data indicates that *TFF1* alleles in cells induced with BPA are maximally active and burst on average once an hour. However, there are fewer active alleles in the population of cells induced with BPA.

### ERα recruitment to *TFF1* and *GREB1* promoters reflects the differential transcriptional responses to BPA

The pioneering factor FOXA1 is one of the main drivers in the establishment of ERα-mediated transcriptome in response to E2 [29–31]. FOXA1 plays a pivotal role in shaping the chromatin structure of E2-ERα target genes in both healthy and cancerous tissues [32, 33]. To determine if recruitment of ERα and FOXA1 was perturbed at the *TFF1* locus after BPA treatment, we profiled their chromatin occupancy using Cleave Under Targets and Tagmentation (CUT&Tag). We treated cells with saturating concentrations of E2, BPA or vehicle for 8h since our single-molecule imaging data suggests the maximum difference in *TFF1* transcription is observed 8h after induction. E2 induced a global increase in ERα binding, with 10,565 called peaks observed and 7,812 called peaks in the vehicle control. Similarly, BPA treatment resulted in increased ERα binding, with 9,058 called peaks. Among the 9,058 ERα binding sites identified following BPA exposure, 4,816 peaks were found to be shared with the E2 induction. This resulted in a total of 19,044 ERα peaks called in our CUT&Tag data set. We focused the rest of our analysis on the 10,565 peaks called in cells induced with E2 (Figure 3A and B). From these results we observe BPA treatment leads to ERα recruitment at the majority of the E2 induced peaks.

**Figure 3:**
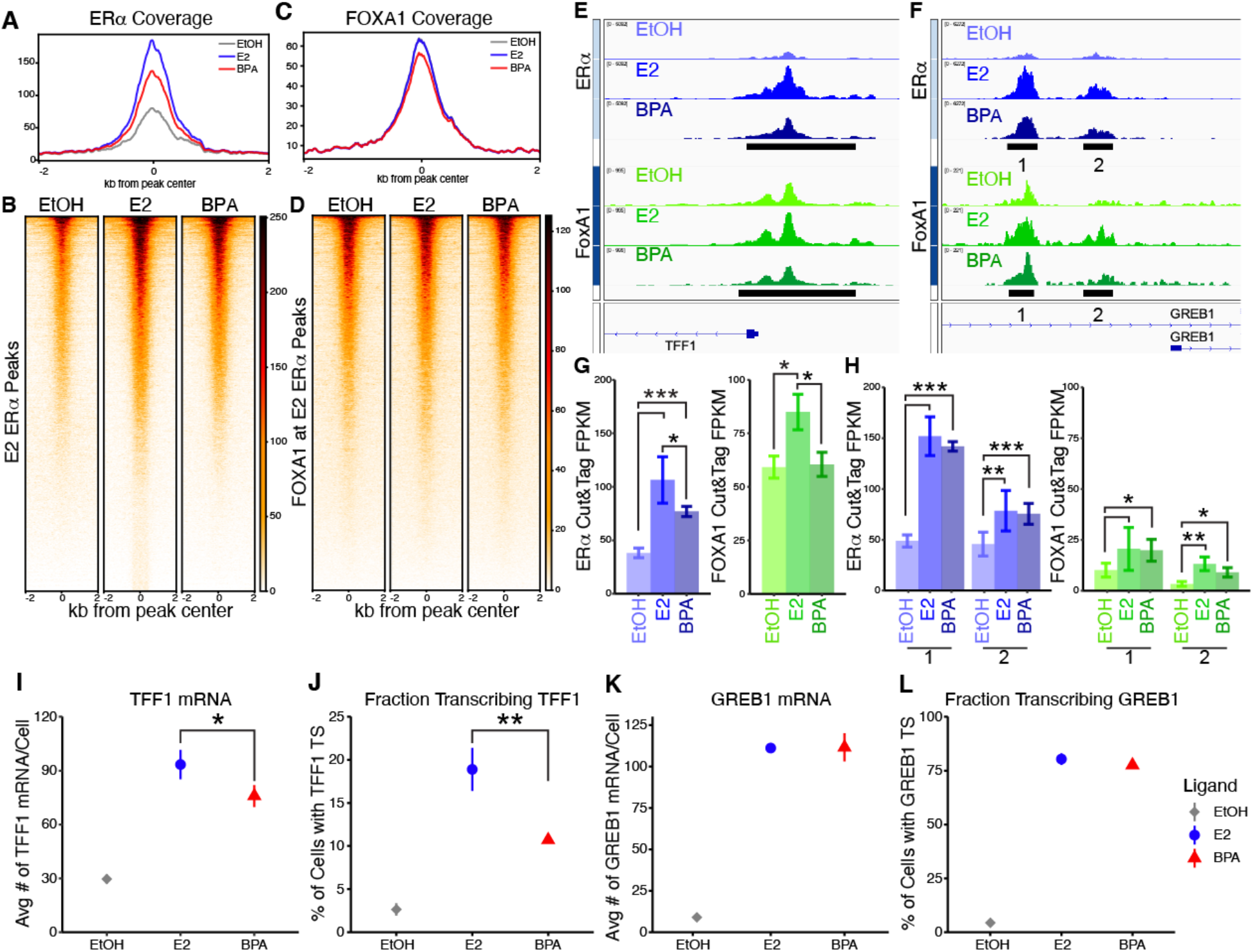
A. Heat map of ERα recruitment at 10,565 ERa peaks 8h after E2. B. Heat map of FOXA1 coverage around E2 ERa peaks. C. Meta profile of ERα and FOXA1 D. CUT&Tag coverage at E2 ERα peaks. P-values were calculated by using the average signal of a 500 bp window centered on the center of the peak and a Wilcoxon Rank Sum/Mann-Whitney test was performed. E. Browser track of the ERa and FOXA1 peaks at the TFF1 promoter. Peaks are annotated by black bars. F. Quantification of ERa and FOXA1 peaks at the TFF1 promoter. P-values determined using DESeq2. G. Browser track of the ERa and FOXA1 peaks at the GREB1 promoter. Peaks are annotated by black bars. H. Quantification of ERa and FOXA1 peaks at the GREB1 promoter. P-values determined using DESeq2. I. smFISH results of *TFF1* mRNA accumulation in cells induced with E2 and BPA. P-values were calculated with t-tests. J. smFISH results of the fraction of cells transcribing *TFF1* in cells induced with E2 and BPA. P-values were calculated with t-tests. K. smFISH results of *GREB1* mRNA accumulation in cells induced with E2 and BPA. P-values were calculated with t-tests. L. smFISH results of the fraction of cells transcribing *GREB1* in cells induced with E2 and BPA. P-values were calculated with t-tests.

We next extracted FOXA1 signal at the 10,565 ERα peaks, however, if a FOXA1 called peak was within 150bp of any of these ERα peaks, we recentered the FOXA1 signal on that FOXA1 peak center (Figure 3C and D). ERα recruitment was significantly reduced by BPA induction relative to E2 (p=3.86e-58, Figure 3A). Additionally, BPA induction resulted in a modest, but significant reduction in FOXA1 recruitment to these sets of peaks (p=0.0435, Figure 3C). To link ERα and FOXA1 recruitment to the fraction of cells transcribing a given estrogen responsive gene we calculate the ratio of ERα recruitment in E2 induced cells to BPA induced cells. ERα recruitment at the *TFF1* promoter was significantly reduced by more than 25% in the presence of BPA, and this reduction was mirrored by a decrease in FOXA1 recruitment (Figure 3E and G). Surprisingly, at the promoter of the estrogen responsive gene *GREB1*, BPA bound ERα recruitment was reduced by less than 10% compared to E2 bound ERα, while FOXA1 binding remained largely unaffected by BPA induction (Figure 3F and H). To determine if ERα and FOXA1 recruitment to these promoters predicted the fraction of actively transcribing cells we designed probes for GREB1 (Sup Fig A) and performed smFISH of both genes. As previously shown, BPA induction significantly reduced *TFF1* mRNA accumulation and reduced the fraction of cells transcribing *TFF1* (Figure 3I and J). However, BPA does not alter *GREB1* mRNA accumulation or the fraction of cells transcribing *GREB1* (Figure 3K and L). These results indicate that ligand-bound ERα and FOXA1 recruitment to gene promoters strongly correlate with the fraction of cells transcribing those estrogen responsive genes.

### BPA results in altered structure at *TFF1*’s promoter

To determine if differential recruitment of ERα and FOXA1 altered open and transcriptionally permissive chromatin, we examined marks of active transcription K27ac and chromatin accessibility with K27ac CUT&Tag and ATAC-seq respectively. Globally, both E2 and BPA treatment resulted in increased K27ac deposition at ERα peaks (p=0.0002 and p=0.003 respectively). We looked specifically at the promoters of *TFF1* and *GREB1* and extracted the reads under the K27ac and ATAC peaks. At the *TFF1* promoter, induction with either E2 or BPA significantly increased K27ac deposition and chromatin accessibility, with no significant difference between treatments (p=0.0003 and 0.0034 respectively, Figure 4A and B). At *GREB1*’s promoter, E2 induction significantly increased K27ac deposition and chromatin accessibility. BPA induction also significantly increased both K27ac and ATAC signals over vehicle control, however, levels of K27ac deposition were reduced compared to E2 induction (Figure 4C and D). This decrease in K27ac in BPA induced cells does not correlate with a reduction in the fraction of cells transcribing *GREB1* (Figure 3L). Our analysis of the K27ac and chromatin accessibility indicates that induction with BPA results in similar levels of open chromatin at the promoter of *TFF1* and contrasts with the reduced fraction of inactive alleles.

**Figure 4:**
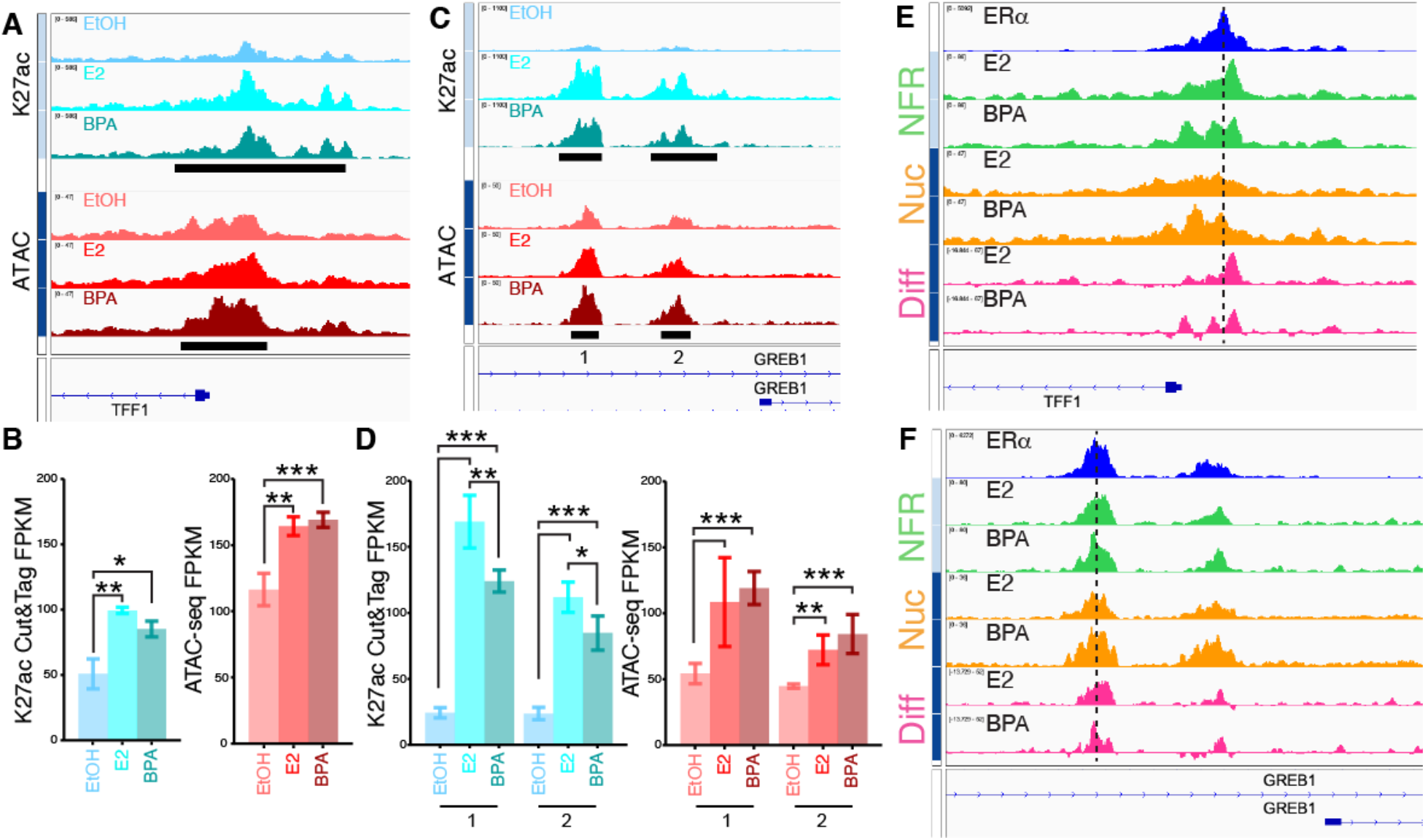
A. Browser track of *TFF1* promoter showing K27ac binding and chromatin accessibility. Peaks are annotated by black bars. B. Quantification of K27ac binding and accessible chromatin peaks at the *TFF1* promoter. P-values determined using DESeq2. C. Browser track of *GREB1* promoter showing K27ac binding and chromatin accessibility. Peaks are annotated by black bars. D. Quantification of K27ac and accessible chromatin peaks at the *GREB1* promoter. P-values determined using DESeq2. E. Browser tracks of ERα peak at the promoter of TFF1 and tracks of ATAC-seq nucleosome free reads (NFR) mononuclesomal reads (Nuc) and the difference of the two previous tracks (Dif). Positive values are enriched in NFR tracks and negative values are enriched in Nuc reads. F. Browser tracks of ERα peak at the promoter of GREB1 and tracks of NFR, Nuc, Dif.

To gain more insight into the structure of the open chromatin at the promoter of *TFF1* we subset the ATAC reads into Nucleosome Free Reads (NFR, <110bp reads) and mononucleosome length reads (150-250bp). In cells induced with BPA we observed a dramatic depletion in NFR reads approximately 160bp and 305bp upstream of *TFF1’s* transcription start site (TSS). We also observe that the mononucleosome reads reveal the converse pattern (Figure 4E). When the mononucleosome reads are subtracted from the NFR reads they reveal more strongly positioned nucleosomes 130bp and 300bp upstream of *TFF1’s* TSS and 85bp downstream of the TSS with BPA treatment (Figure 4E). Interestingly, one of the nucleosomes with increased positioning in BPA induced cells falls directly under the CUT&Tag ERα peak proximal to *TFF1’s* promoter. At the promoter of *GREB1,* BPA induction resulted in moderate alterations to chromatin structure, including differences in the NFR reads and a slight increase in nucleosome positioning at the proximal ERα peak. However, the *GREB1* promoter structure was already phasic, whereas *TFF1* was considerably remodeled into a phasic structure. Taken together these results indicate that BPA induction results in increased open chromatin at ERα target genes, but a subset of genes, including *TFF1*, have altered promoter structure that correlates with a reduction in the fraction of actively transcribing cells.

### BPA bound ERα disrupts the enhancer-promoter interaction at *TFF1* locus through cofactor recruitment

Our results suggest that *TFF1* requires recruitment of specific cofactors to properly remodel the promoter. We assessed the impact of two established transcriptional coactivators of ERα, SRC-3 and MED1, on *TFF1* bursting. We employed SRC-3 (siSRC-3) specific or non-targeting (siNT) siRNA pools to reduce SRC-3 levels in hormone-depleted cells which were subsequently induced with saturating levels of E2, BPA, or a vehicle control. We quantified the fraction of cells actively transcribing *TFF1* using smFISH. For the siNT transfected cells we observed the expected significant increase of fraction of actively transcribing cells in the presence of E2 and a significant decrease, relative to E2, when the cells are induced with BPA. Furthermore, when SRC-3 was knocked down, we noted a statistically significant reduction in the basal levels of actively transcribing cells in the vehicle control (p=0.018). Additionally, in siSRC-3 transfected cells, both E2- and BPA-mediated activation of *TFF1* transcription was attenuated and comparable with the siNT control (p=0.535 and 0.983 respectively, Figure 5A). These findings suggest that SRC-3 is responsible for keeping alleles in a transcriptional permissive state in the absence of ligand or residual E2 mediated activity.

**Figure 5:**
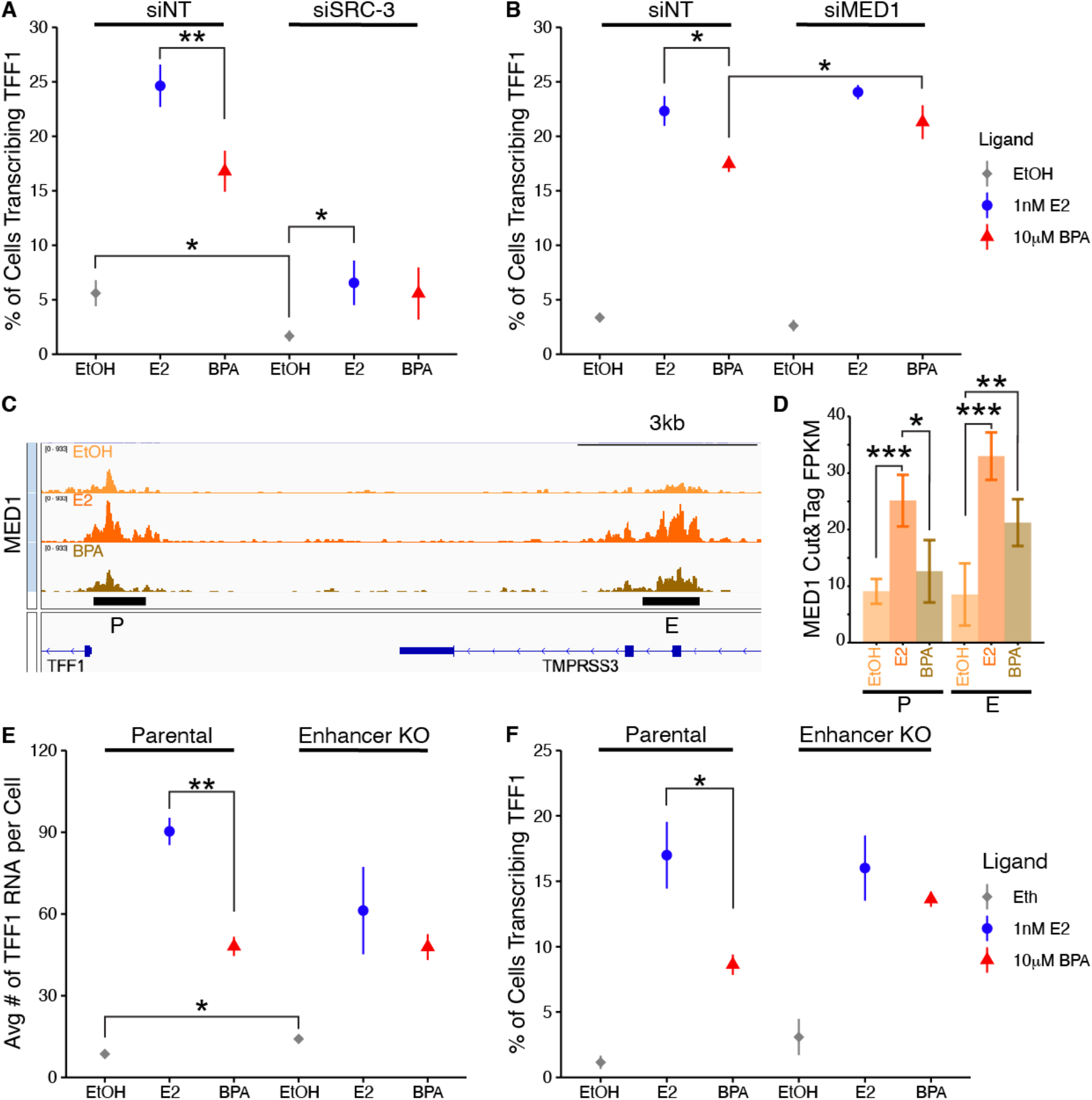
A. smFISH results of the fraction of cells actively transcribing *TFF1* in siNT or siSRC-3 transfected cells induced with E2, BPA, or vehicle control. P-values were calculated with t-tests. B. smFISH results of the fraction of cells actively transcribing *TFF1* in siNT or siMED1 transfected cells induced with E2, BPA, or vehicle control. P-values were calculated with t-tests. C. Browser track of MED1 binding at the *TFF1* locus. Peaks are annotated by black bars. D. Quantification of MED1 peaks at the *TFF1* locus. P-values determined using DESeq2. E. smFISH results of the fraction of cells actively transcribing *TFF1* in parental or *TFF1* enhancer deleted MCF-7 cells induced with E2, BPA, or vehicle control. P-values were calculated with t-tests. F. smFISH results of *TFF1* mRNA accumulation in parental or *TFF1* enhancer deleted MCF-7 cells induced with E2, BPA, or vehicle control. P-values were calculated with t-tests.

To determine how MED1 impacted the fraction of actively transcribing cells with BPA treatment, we quantified cells with *TFF1* transcription sites using smFISH in MED1-knockdown cells induced with saturating levels of E2, BPA, or vehicle. We observed similar ratios of actively transcribing cells when siNT was transfected, consistent with our previous results. Knockdown of MED1 in the presence of E2 does not elicit any change in the fractional response of cells (Figure 5B). Interestingly, knockdown MED1 partially rescued the BPA-mediated reduction in transcriptional response without affecting the basal levels of *TFF1* transcription in the vehicle control (Figure 5B). We next investigated MED1 occupancy at the *TFF1* locus to determine if BPA treatment induces changes in recruitment at the endogenous gene locus using CUT&Tag (Figure 5C and D). MED1 recruitment increases at both the *TFF1* promoter and enhancer when cells are treated with E2 or BPA. However, while MED1 recruitment remains unchanged by BPA at the enhancer region, there is a significant decrease at the *TFF1* promoter (Figure 5D). Since ERα recruitment also decreases at the promoter and does not significantly change at the enhancer, our data suggests a disruption in the communication between the promoter and enhancer.

Given the Mediator complex’s role in regulating promoter and enhancer interactions we next asked how removal of *TFF1’s* proximal enhancer, located less than 10 kb upstream of the promoter, affected the fraction of cells transcribing *TFF1* in BPA induced cells. We used a previously described cell line where the proximal enhancer region was deleted in MCF-7 cells [10]. Following induction with saturating levels of E2, BPA, or vehicle, we quantified mRNA accumulation per cell and the fraction of actively transcribing cells using smFISH. In the enhancer deletion cells (Enhancer KO), *TFF1* mRNA accumulation induced by E2 is reduced. BPA induction resulted in a similar level of *TFF1* mRNA accumulation as E2 treatment (Figure 5E). Intriguingly, the deletion of the enhancer leads to a similar fraction of actively transcribing cells treated with BPA, resulting in a level of activity similar to that of E2-induced cells (Figure 5F). In summary, our findings from MED1 knockdown and enhancer deletion results suggest that BPA disrupts the coordination between the *TFF1* promoter and enhancer, limiting the ability of the BPA-ERα complex to activate *TFF1* alleles. Consequently, BPA bound ERα primarily induces transcription from alleles that are already in an active state.

### BPA bound ERα is a limiting factor for fractional response in a subset of E2 responsive genes

To identify other genes with altered expression in response to BPA we performed RNA-seq of cells that had been induced with saturating levels of E2 or BPA for 24h. We found 964 significantly up regulated genes and 1,046 significantly down regulated genes (Figure 6A) in cells induced with E2. Similarly, BPA treated cells had a robust response with 831 significantly up regulated genes and 837 down regulated genes (Figure 6B). As expected *TFF1* was significantly upregulated in both RNA-seq datasets, thus we focused on the 964 genes that were upregulated by E2 for further analysis. Of the 964 genes up regulated by E2, 33.5% of the genes were significantly decreased in cells treated with BPA (Figure 6C, purple). *TFF1* did not pass our FDR cut off of 0.05. However, repeated RNA FISH experiments have shown that BPA induction significantly reduces *TFF1* mRNA accumulation, and likely reflects differences in sensitivity between the techniques. We identified 20 genes with expression levels reduced by at least 30% relative to E2 including *EGR3*. To determine if the fraction of cells actively transcribing *EGR3* was altered by BPA we designed FISH probes for *EGR3* (Sup Fig B). We observed a significant reduction in *EGR3* mRNA (Figure 6D) and a significant reduction in the fraction of cells actively transcribing *EGR3* in cells induced with BPA (Figure 6E). This analysis indicates that a majority of E2 responsive genes are similarly induced by BPA. However, our data suggests the genes reduced RNA output by BPA have a smaller fraction of active alleles.

**Figure 6:**
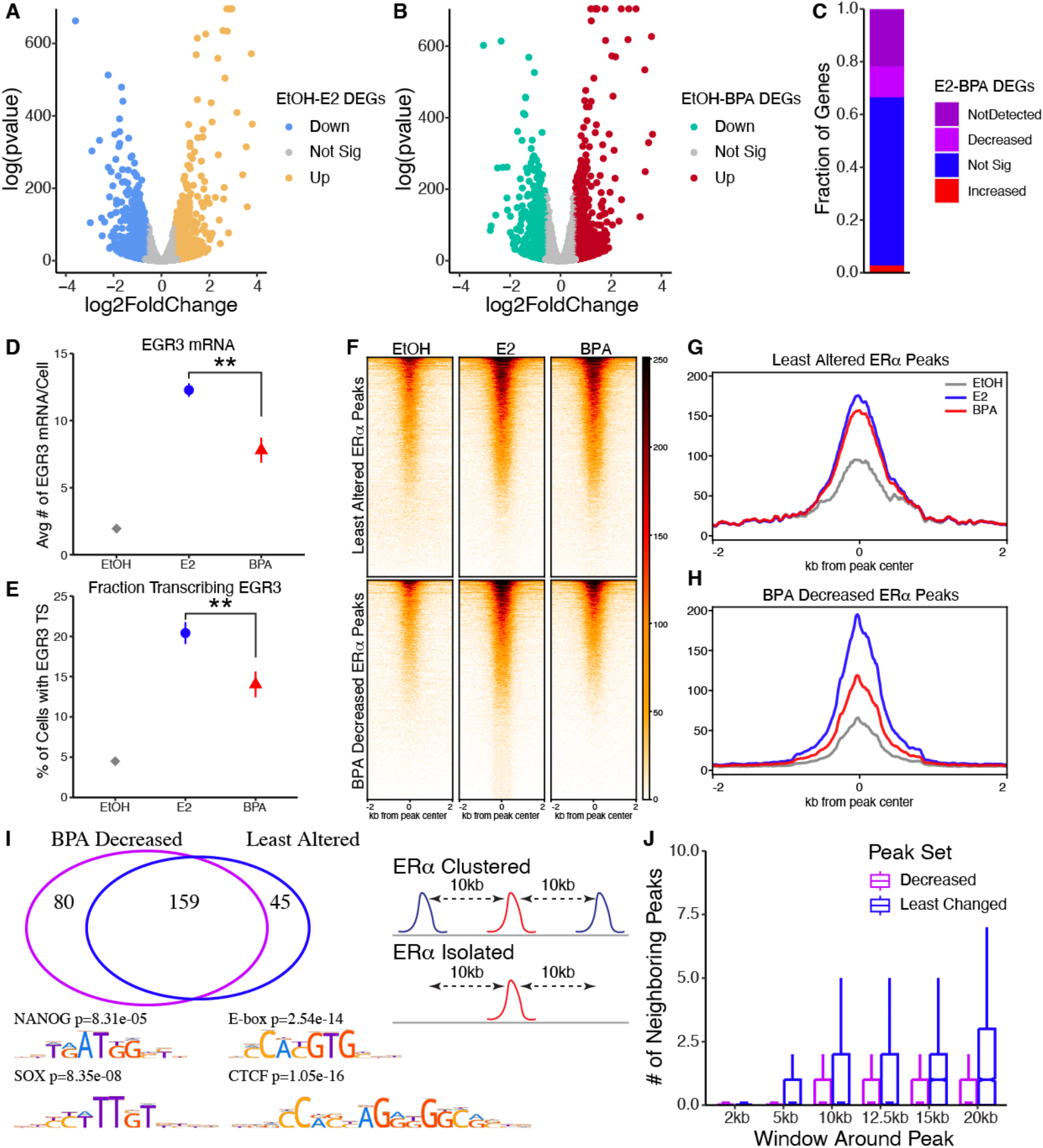
A. Volcano plot of the log2 fold change vs -log(pvalue) of E2 induced gene expression changes. Genes with a padj <0.05 and a fold change of 1.5 or more were considered significantly changed, down regulated genes in blue and upregulated genes in yellow. B. Volcano plot of the log2 fold change vs -log(pvalue) of BPA induced gene expression changes. Genes with a padj <0.05 and a fold change of 1.5 or more were considered significantly changed down regulated genes in green and up regulated genes in red. C. A bar plot of the E2 upregulated genes. Purple indicated the genes decreased by BPA with a padj <0.05, red indicated the genes increased by BPA relative to E2 with a padj <0.05. D. smFISH results of *EGR3* mRNA accumulation induced with E2, BPA, or vehicle. P-values were calculated with t-tests. E. smFISH results of the fraction of cells actively transcribing *EGR3* induced with E2, BPA, or vehicle. P-values were calculated with t-tests. F. Heat map Least Altered and BPA Decreased ERα peaks. G. Meta profile of Least Altered ERα peaks. H. Meta profile of BPA Decrease ERα peaks. I. Venn diagram of the unique motif identified in the ERα peak sets by MEM. J. Comparison of the number of neighboring peaks within a given window length in Least Altered and Decreased peak set. The representation of Clustered or Isolated ERα peaks.

To determine if differential ERα recruitment reflects the fractional response, we calculated the ratio of reads under ERα peaks in E2 induced cells to BPA treated cells. We sorted ERα peaks by this ratio and split the peak-set in half (Figure 6F-H). As expected, the promoter of *TFF1* and the enhancer peaks of *EGR3* were in the BPA Decreased half and *GREB1* was in the Least Altered half. Next, we performed motif enrichment on the peak sets. As expected, common motifs between the two sets included ESR1 and FOXA1. Interestingly, the least altered peaks were enriched in E-box transcription factors including MYC, MAX and CTCF. The most altered peaks were enriched for stem cell related transcription factors such as SOX and NANOG (Figure 6I). Finally, given that there were multiple peaks at the promoter of *GREB1* while *TFF1’s* promoter peak was relatively isolated, we asked if the least altered ERα peaks were more likely to exist in clusters. Using a window around the peak, we observed that the least altered ERα peaks have more neighboring peaks than the most altered ERα peaks (Figure 6H). Our data suggest that clustered peaks are more resistant to the effects of BPA treatment compared to more isolated ERα peaks and are enriched for CTCF that stabilize enhancer promoter interactions.

## Discussion

We have shown that the complexes recruited by E2 bound ERα serve two functions. They switch *TFF1* alleles from a transcriptionally nonpermissive state to a permissive state and they activate alleles already in the permissive state (Figure 7, step 2). We find that BPA treatment perturbs this response, BPA bound ERα is unable to transition alleles into the permissive state (Figure 7). However, at some alleles BPA can maximally induce transcriptional bursts with the same initiation rates as E2, approximately once an hour. We show that ERα recruitment reflects the fractional response, and this reduced binding coincides with reduced cofactor recruitment. Despite this reduced recruitment, BPA treated cells can increase accessibility at the *TFF1* promoter. However, the promoter is in an altered chromatin structure (Figure 7). These data strongly indicate that the transition from the nonpermissive state to permissive state requires the involvement of more regulatory elements including cofactors such as FOXA1.

**Figure 7:**
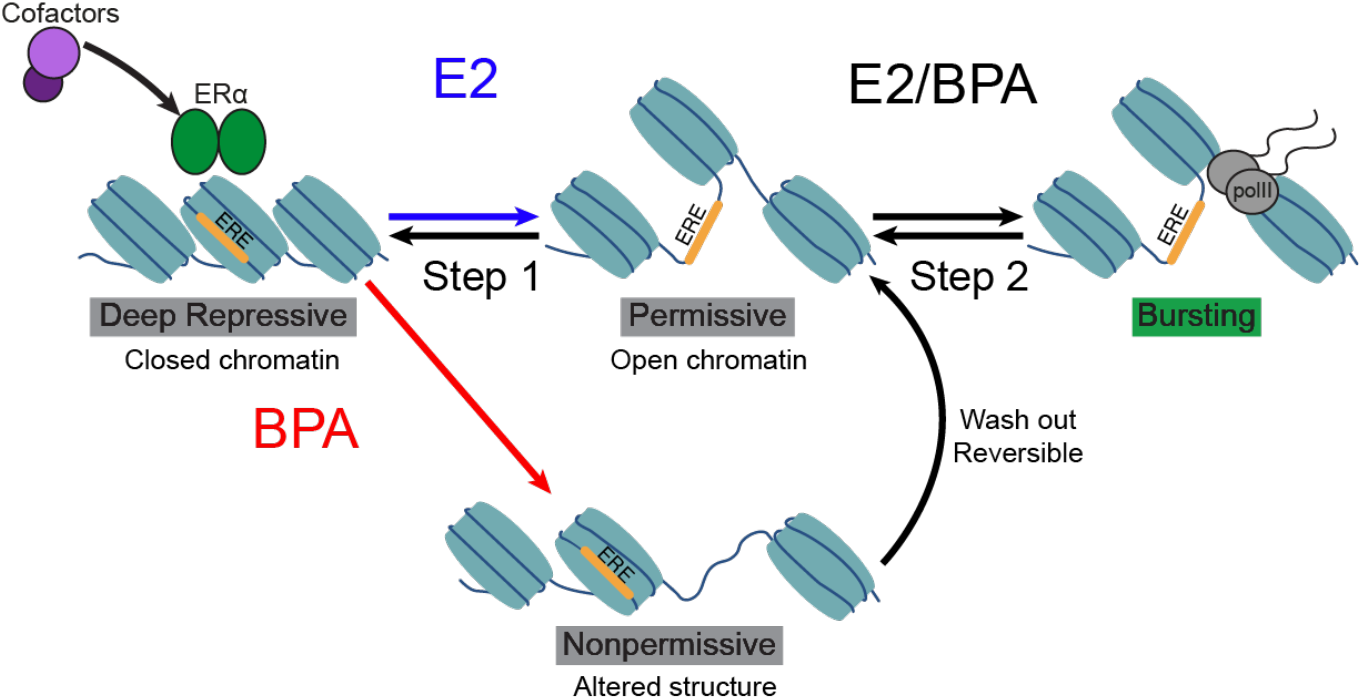
A model representing the predominate transcriptional states governing transcription of *TFF1*. Arrows indicate state transitions.

Our study establishes a correlation between the fraction of cells actively transcribing *TFF1* and *GREB1,* and the recruitment of ERα and FOXA1 at the gene loci. This result, in the context of studies which profiled the protein-protein interactions formed by ERα in the presence of E2 and BPA [20, 21], suggest that activation of *GREB1* does not depend on the same cofactors as *TFF1*. Supporting this speculation, different subsets of cofactors are necessary for maximal activation of E2 responsive genes [34]. Collectively, these results suggest that a subgroup of genes exhibit a greater dependence on the composition of the complex recruited by ERα for activating alleles from a nonpermissive state.

Our work shows that these transcriptional state switches are not primarily governed by the accumulation of K27ac or open chromatin. Induction with both E2 and BPA results in significant increases of K27ac and open chromatin at the promoters of both *TFF1* and *GREB1*. Specifically, at *TFF1* we did not observe a statistical difference between conditions in either K27ac or open chromatin. However, our ATAC data indicates that in cells induced with BPA, the promoter of *TFF1* exhibits more open chromatin flanking the ERα peak summit. Furthermore, two nucleosomes flanking the TSS of *TFF1* display altered positioning, suggesting an altered footprint of the complex recruited by ERα. This observation and our imaging data suggests a new transcriptionally nonpermissive state induced by BPA (Figure 7, altered structure). When BPA is washed out and cells are induced with E2, the fraction of cells transcribing *TFF1* recovers to the same level of cells induced with only E2. These results lead us to conclude that ERα bound by BPA lacks the cofactors necessary to effectively remodel chromatin to transcriptionally permissive states at specific genes.

We focused on two cofactors, SRC-3 and MED1, shown to interact with ERα in the presence of E2 and BPS, but not BPA. Previous work has shown that these cofactors are critical to the estrogen response and are required for the switch from acute to sustained activation of E2 target genes [35]. We observe that when cells are induced with E2 and BPA, the fraction of cells with a *TFF1* transcription site continues increasing for E2 induced cells, but plateaus for cells treated with BPA. When the model is considered with our observations, we favor a revised model where the initial estrogen response occurs at alleles that are in transcriptionally permissive states, and the sustained activation arises from alleles transitioning from closed chromatin to permissive states.

We show that MED1 recruitment is reduced at the *TFF1* locus in the presence of BPA, consistent with prior observations indicating that ERα-MED1 interaction is compromised with BPA. Importantly, deletion of the enhancer of *TFF1* results in a similar fraction of cells actively transcribing *TFF1*. These results indicate that transcription of *TFF1* is reduced by BPA through reduced cooperation between the enhancer and promoter of *TFF1*. This further emphasized the critical role of the proper combination of cofactors in productive enhancer and promoter interactions. Our data also suggests that this impediment can be overcome by a higher local concentration of ERα binding sites. In summary, our result show that for a subset of genes, ERα requires the appropriate composition of cofactors to activate alleles that are in non-permissive states. However, for alleles that are in a permissive state, the reduced complex recruited by ERα suffices to maximally activate alleles. Moreover, a subset of genes, ERα can overcome alterations in the complex recruited by ERα, resulting in full activation by BPA. These results shed light on the mechanisms of transcriptional state switching and has implication for the regulation of environmentally sensitive genes.

## Methods

### Cell Culture

MCF-7 cells (ATCC) were grown in MEM supplemented with 10% FBS (Sigma), 2mM Glutamine and 1X Penstrep (Gibco). smFISH experiments were performed in 12 well plates using cells plated on 18mm No. 1.5 coverslips. Cells were plated on coverslips and allowed to recover for 2 days post plating before being hormone depleted. Cells were hormone depleted by washing with Phenol free MEM supplemented with 10% Charcoal/Dextran Treated FBS (Sigma), 2mM Glutamine and 1X Penstrep. Cells were returned to the incubator for 1h. This step was performed again. This procedure for hormone depletion was followed for all subsequent experiments. After 2 days of hormone depletion, cells were induced with ligand. For the dose response and steady state experiments, cells were induced for 65h, other experiments were performed as described.

For live cell *TFF1-MS2* imaging MCF-7 cells were plated on glass bottom (Nunc) dishes and allowed to recover for multiple days prior to hormone depletion. Cells were hormone depleted for 2 days prior to addition of ligand 8 hours before imaging.

For CUT&Tag and ATAC-seq MCF-7 cells were plated in 10cm dishes and allowed to recover for 2 days. Cells were subsequently hormone depleted for 2 days prior to induction with ligand. 8 hours after induction with ligand nuclei were isolated for CUT&Tag or ATAC-seq.

Cells were plated for washout experiments and allowed to recover for 2 days. Cells were then hormone depleted for 2 days prior to induction with the primary ligand for 8h. Cells were then briefly rinsed with hormone depleted media before induction with the final ligand for 4h.

siRNA experiments were performed by allowing cells to recover for 2 days after plating before hormone depletion. 24h after hormone depletion cells were transfected with siRNA smart pools (Dharmacon) using Lipofectamine RNAiMAX Transfection Reagent and the manufacturers protocol. 24h after transfection cells were induced with ligand for an additional 24h prior to fixation or harvesting for other analysis.

### smFISH

The *TFF1* exon probe set (16 probes) and intron probe set (79 probes) were previously described [10]. The *GREB1* intron and exon probe sets (48 probes each) and *EGR3* exon probe set (45 probes) were designed using Stellaris Probe Designer (https://www.biosearchtech.com/stellaris-designer) using a masking level of 5, oligo length of 20, minimum spacing of 2 nucleotides. *TFF1* and *GREB1* exon probe sets were ordered from Biosearch Stellaris in Quasar 570. *TFF1* and *GREB1* intron and *EGR3* exon probe sets were ordered from Biosearch Stellaris in Quasar 670. 3 hybridization chain reaction (HCR) doublets were designed to the intron of *EGR3*. Briefly, Primer3 was used to generate candidate probes. HCR initiator doublets, probes exactly 2bp apart, were selected using a custom python script. HCR initiator probes and fluorescently labeled hairpins were ordered from Integrated DNA Technologies (IDT).

We used the Stellaris smFISH protocol as previously described [10]. Briefly, cells were fixed with 4% PFA for 10 min, washed with PBS, then permeabilized with 70% ethanol overnight at 4°C. Probe hybridization at 37C was performed following the adherent mammalian cell Stellaris RNA FISH Protocol. Finally, coverslips were mounted using Prolong Gold with DAPI and allowed to dry shielded from light for 24h before imaging.

For smFISH using HCR probe sets sequential HCR and Stellaris smFISH was performed as previously described [36]. Briefly, HCR FISH was performed first according to the protocol described in [37] with minor modifications. Probe wash buffer and hybridization buffers both lacked citric acid. Initiator probes were hybridized over night at 37°C and signal amplification was performed at room temperature for 90 min. Following amplification cells were washed 5 times in 5x SSCT before proceeding to the adherent mammalian cell Stellaris RNA FISH Protocol described above.

### Microscopy

smFISH images were acquired using a custom microscope built on an ASI (www.asiimaging.com/) Rapid Automated Modular Microscope System (RAMM) base fitted with an ASI MS-2000 Small XY stage, a Hamamatsu ORCA-Flash4 V3 CMOS camera (https://www.hamamatsu.com/, C13440-20CU), an ASI High Speed Filter Wheel (FW-1000), ASI MS-2000 Small XY stage, and a Zeiss C-Apochromat 40x / 1.20 NA UV-VIS-IR objective. DAPI, Quasar 570, Quasar 670, or Cy5 were excited using a Lumencore SpectraX (https://lumencor.com/) with violet, green, and red filters. The microscope was controlled using Micro-Manager (Edelstein et al., 2010) and twenty-five fields of view consisting of a 10-micron Z stack with 0.5 micron intervals.

Live cell imaging was performed using a Andor Dragonfly Spinning Disk Confocal (https://andor.oxinst.com/) using 37°C incubation and 5% C02. Images were acquired using 488nm excitation, on a 60x objective with a pinhole size of 40μm. 7 z planes were acquired with 0.8 micron interval every 100s for 512 frames totaling 14.2h. Analysis was performed using maximum intensity projections.

### Image analysis

Analysis of smFISH data was performed on maximum intensity projections using a custom python script. Briefly smFISH spots were called by fitting a 2D Gaussian mask and performing local background subtraction. Masks of the cytoplasm and nuclei were generated using Cellprofiler [38]. Transcriptions sites (TS) were identified in the nuclear mask by intron and exon smFISH spots with a Euclidean distance less than or equal to 5 pixels. Number of nascent transcripts was determined on a per cell basis by extracting the intensity of the exon signal at the identified TS and dividing by median intensity of at least 5 exon smFISH spots detected in the cytoplasmic mask.

Analysis of *TFF1-MS2* active and inactive durations was determined using custom IDL software, Localize, as previously described [10, 39]. Briefly, individual nuclei were tracked over all 512 frames using TrackMate [40] and the center coordinates of each nuclei was passed to a custom python script which generated a cropped time series for each nuclei. These cropped time series were used by Localize to segment active TSs by fitting a 2D Gaussian mask and performing local background subtraction. Traces were generated by using the track function in Localize which extracts the TS intensity and the background intensity for each frame in between called TS. Tracks were manually inspected to ensure different alleles in the same nuclei were not being tracked. Tracks were then analyzed in python by a Hidden Markov Model (HMM) algorithm to detect active TS and converted to a binary trace denoting frames where *TFF1-MS2* as active and inactive. Finally, cumulative distributions of active and inactive durations were compiled from many traces.

The fraction of active cells was calculated by calling active TS using deepBlink [41] to analyze each cropped nuclei time series extracted using TrackMate. For each cell the first frame where a TS was detected using deepBlink was recorded, and the cell was considered active. The fraction of active cells was determined by dividing the number of active cells by the total number of tracked cells for each individual frame.

### CUT&Tag

CUT&Tag was performed following the Bench top CUT&Tag V.3 protocol [42]. Briefly, we used 100K nuclei isolated from E2, BPA, or vehicle treated MCF7 cells per reaction. 10 µl BioMag-Plus Concanavalin (Bangs Laboratories) were used per reaction to immobilize the nuclei. Then we added 1: 50 mouse ERα (sc-8005, Santa Cruz), 1:25 rabbit FOXA1 (#53528, Cell Signaling Technology), 1:50 mouse H3K27ac (#MA5-23516, Invitrogen), or 1:25 rabbit MED1 (#51613, Cell Signaling Technology) antibody and incubated for 2 hours. After one hour of secondary antibody incubation 1.25 µl of CUTANA pAG-Tn5 adapter complex was used to load the enzyme to the antibody bound regions. One hour of tagmentation at 37°C was followed by DNA extraction using MinElute PCR Purification Kit (Qiagen). Extracted DNA was subjected to PCR amplification using unique primers sets (Nextera XT v2 Full set (N7-S5)). Number of PCR cycles were determined specifically to prevent overamplification. We amplified ERα, FOXA1, H3K27ac, or MED1 with 14, 15, or 17 PCR cycles, respectively. Libraries were size selected using AMPure XP Beads (Beckman Coulter) was with 1.3X ratio. We quantified and assessed the quality of the libraries with Qubit Flex and Tapestation and then sequenced the libraries with 50 bp paired-end reads on Illumina high output NovaSeq SP.

### ATAC-seq

ATAC-seq samples were prepared following the protocol described in [43] with minor modifications. First 50K nuclei were isolated in technical duplicate per sample. DNA was tagmented for 30 min at 37C using the Illumina Tagment DNA kit (20034211). Tagmented DNA was cleaned up using the MinElute PCR Purification Kit (Qiagen). The number of PCR cycles used for library amplification was determined by qPCR. Finally, a double-sided size selection was performed using AMPure XP Beads (Beckman Coulter). Libraries quality was assessed using by Tapestation and the technical duplicate of each sample with the most pronounced nucleosome banding was sequenced.

### CUT&Tag and ATAC-seq Analysis

Briefly, CUT&Tag and ATAC-seq reads were processed using pipelined adapted from [44]. First adaptors were trimmed using Cutadapt and reads were aligned to the hg38 genome assembly using Bowtie2. Samtools was used to sort the reads and deduplication was performed using Picard Tools (http://broadinstitute.github.io/picard). Samtools was also used to separate reads into read lengths of 1-110 bp, nucleosome free, and 150-250 bp, mononucleosome reads. Peaks were called using MACS2 and filtered to keep peaks that were called in more than one replicate of at least one condition. Peaks were quantified with the featureCounts tool and differential analyses was performed using the DESeq2 R package. Motif analysis was run by using the bed coordinates in the SEA tool in the MEM suite. Window analysis was performed using the count function in bedtools window.

### RNA-Seq

Cells were induced with E2, BPA, or vehicle for 24 hours and then harvested. Total RNA was isolated using RNeasy Mini Kit (Qiagen). The quantity and quality of the RNA samples were assessed by Tapestation. Polyadenylated mRNAs were selected using Dynabeads Oligo(dT)25 (Invitrogen #61005) and libraries were prepared with NEBNext Ultra Directional RNA Library Prep Kit for Illumina (NEB #E7760) following manufacturer’s instruction. Libraries were size selected using AMPure XP Beads (Beckman Coulter). Libraries were quantified Qubit Flex and Tapestation and then sequenced with 75 bp single-end reads on Illumina NextSeq high-output.

Reads were filtered so that only those with a mean quality score of 20 or greater were kept. Adapter was trimmed using Cutadapt version 3.7. Reads were aligned to the hg38 genome assembly using STAR version 2.6.0c. Counts were obtained using the featureCounts tool from the Subread package version 1.5.1 with the GENCODE basic gene annotation version 44. Differential expression was quantified using the DESeq2 R package version 1.34.0.

## Acknowledgements

This research was supported in part by the Intramural Research Program of the NIH. The Fluorescence Microscopy and Imaging Center (NIEHS), specifically Jeff Tucker and Erica Scappini, for use of the Andor Dragonfly Spinning Disk Confocal/HILO Microscope. The Epigenomics and DNA Sequencing Core (NIEHS) Jason Malphurs and Brian Papas for sequencing and initial QC of CUT&TAG/ATAC-seq libraries. Krystal Orlando and Yang Liu (Wade Lab ESCBL) for trouble shooting CUT&TAG/ATAC-seq with us. Jackson Hoffman (Archer Lab ESCBL) for his support and encouragement on this project.

## Supplemental figures

**Figure S1:**
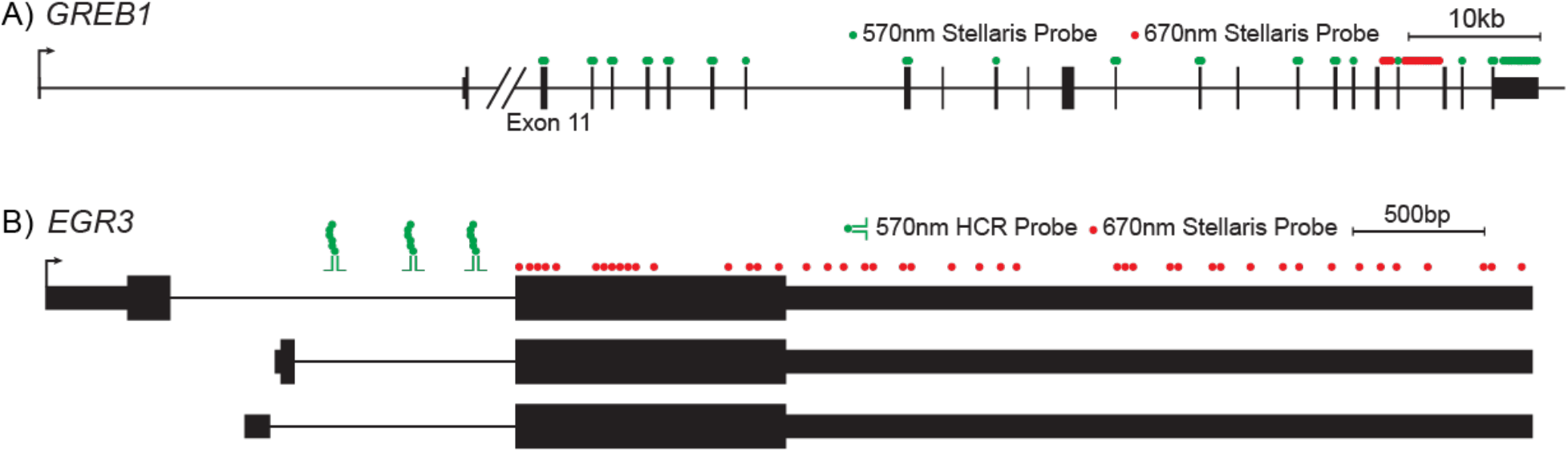

